# The rate of false polymorphisms introduced when imputing genotypes from global imputation panels

**DOI:** 10.1101/080770

**Authors:** Ida Surakka, Antti-Pekka Sarin, Sanni E Ruotsalainen, Richard Durbin, Veikko Salomaa, Mark J Daly, Aarno Palotie, Samuli Ripatti, SISu project group

## Abstract

Previous studies^1,2^ have shown that large multi-population imputation reference panels increases the number of well-imputed variants. However, to our knowledge, no previous studies have evaluated the rate of introduced variation in monomorphic sites of the study population when using imputation panels with admixed populations. In this study we evaluate the rate of false positive variants introduced by the imputation of Finnish genotype data using global reference panels (Haplotype Reference Consortium^1^; HRC, and the 1000Genomes project Phase I^3^; 1000G) and compare the results to a Finnish population-specific reference panel combining whole genome and exome sequenced samples. In sites that were monomorphic in our test set, we observed high false positive rates for the global reference panels (4.0% for 1000G and 2.6% for HRC) compared to the Finnish panel (0.26%). This rate was even higher (7.4%) when using a combination panel of 1000G and Finnish whole genome sequences with cross-panel imputation.

---

Genotype imputation is a method where a study set of observed microarray based genotypes is extended using a set of sequenced or densely genotyped haplotypes from a set of reference haplotypes. Besides the quality and source of study genotypes, one of the crucial factors in the imputation precision is the size and quality of the reference panel. Global imputation panels have become a standard sources of reference haplotypes with the panels from 1000 Genomes Project^3^, and more recently Haplotype Reference Consortium^1^ widely used to improve the genomic coverage given by the microarray data. It has been shown that the number of successfully imputed markers grows as the panel size grows, even with haplotypes from varying origin in the panel^1,2^.

While the large population-specific panels typically allow for high quality imputation^4,5,6^, the combination of population-specific panel and the global panel provides the largest number of successfully imputed variants^6,7^. The larger number of reference haplotypes raises the proportion of high quality imputed variants, particularly in low minor allele frequencies.

Currently, there is also a large number of whole exome sequencing (WES) studies being carried out around the world. A good example is the recently published ExaC database^8^ combining WES data from multiple populations. It is unclear, how much these large-scale whole exome resources contribute to the imputation quality of coding variants.

The focus of genetic association studies has shifted from common genetic variants (minor allele frequency, MAF > 5%) to rare-and low frequency variants that are more challenging to impute accurately because of low number of carrier haplotypes in the reference panels. In case of Finnish population, a founder population with multiple bottleneck effects and internal migration^9^, the population history has caused enrichment of specific rare and low frequency variants in the population.^10^ Similarly many variants seen in Europeans are not present in the Finnish population. The other implication of the Finnish population structure is that there is extended linkage disequilibrium^11^ (LD) compared to more mixed populations and this might facilitate the imputation in Finns in general. These population features have motivated sequencing of high number of Finns to study the downstream health effects of the enriched variants^10^.

The aim of this study is to compare the performance of population specific panel with the whole exome sequence based reference panel extension to the performance of the large-scale global reference panels. This will imply whether usage of the population specific panel instead of the already existing large reference panels is encouraged in the imputation of founder population genotype data.

We used different reference panels or reference panel combinations (all listed in **Table 1**) to test which one performs best in terms of imputation accuracy, false positive rate and number of variants that can be considered to be imputed with good quality using chromosome 17 as an example. The panels we used to evaluate the performance were: 1) 1000 genomes^3^ (1000G) September 2013 release 2) Haplotype Reference Consortium^1^ (HRC) release 1 and 3) a local panel combining 1,941 Finnish low-pass WGS and 4,932 Finnish high-pass WES (http://www.sisuproject.fi). Finns were present also in the global panels with the number in the 1000G being 93 and in the HRC panel 1,941 (the same Finnish low-pass WGS samples are included in the Haplotype Reference Consortium). Before imputation we masked all variation with MAF < 1% for the imputation accuracy assessment.

**Table 1:** Description of the reference panels and tested combinations.

| Reference panel | Number of autosomal variants | Number of individuals |
| --- | --- | --- |
| 1000G | 37,878,821 | 1092 |
| LP-WGS | 13,625,231 | 1941 |
| 1000G + LP-WGS | 41,369,683 | 3033 |
| HRC | 39,235,157 | 32488 |
| WES | 222,364 | 4932 |
| LP-WGS + WES (Local panel) | 13,714,270 | 6873 |

1000G: 1000 Genomes panel; LP-WGS: Finnish low-pass whole genome sequences; 1000G + LP-WGS: Finnish whole genome sequence panel combined with 1000G panel; HRC: Haplotype reference Consortium panel; WES: Finnish high-pass whole exome sequences.

We tested the performance of these panels using a sample set of 10,489 individuals sampled from the National Finrisk Studies^12^. The overall imputation quality, measured as the information score of IMPUTE2 software^13^, of the tested reference panels is compared in the **Figure 1A**. In the comparison, 1000G had more poorly imputed variants (high peak around zero) than well-imputed variants (high peak around one) whereas using HRC panel the proportion of well-imputed (info > 0.7) variants grew notably. Moreover, as can be seen from the figure, the density of the IMPUTE2 information scores was clearly shifting towards one when using the local reference panel consisting of Finnish samples compared to either of the global reference panels.

**Figure 1.**
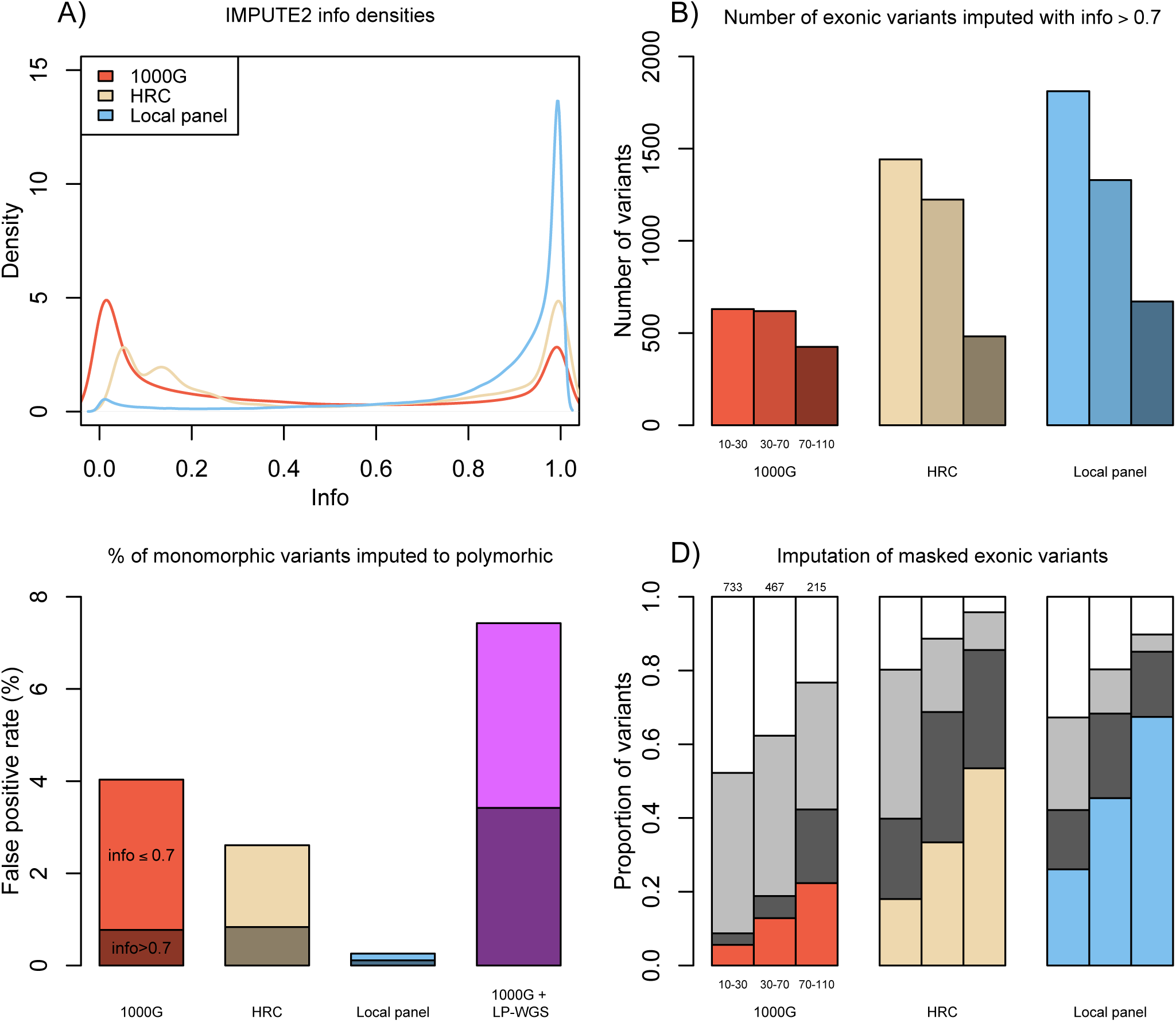
Summary of the imputation results. Panel A) represents the overall imputation quality of the chromosome 17. Panel B) shows the number of well-imputed (info > 0.7) exonic variation with three different minor allele count (MAC) ranges (10-30, 30-70, 70-110) for each of the tested panels. Panel C) illustrates the false positive rate of each panel with division of markers into info ≤ 0.7 and info > 0.7. Panel D) shows the imputation results for the masked variation. Each of the masked variants falls into one of four groups for each of the panels; white = not covered by the panel, light grey = info < 0.7, dark grey = concordance < 0.8, colored = concordance > 0.8. 1000G: 1000 Genomes panel (N = 1,092); HRC: Haplotype reference Consortium panel (N = 32,488); Local panel: Combined Finnish low-pass whole genome sequence and high-pass whole exome sequence panel (N = 6,873); 1000G + LP-WGS: Finnish whole genome sequence panel combined with 1000G panel (N = 3,033).

When looking only at the rare exonic variation, the total number of well-imputed (info > 0.7) variants, with minor allele count (MAC) between 10 and 110 (comparable to a minor allele frequency between 0.05% and 0.5%), was higher when using HRC panel instead of the 1000G (**Figure 1B**). The gain was largest for the lowest MAC range (10-30) where we saw 129% increase, illustrating the effect of reference panel size and/or the number of matching reference haplotypes in imputation as 1000G panel has 93 Finnish samples and HRC has 1,941. However, when comparing the local panel to the biggest panel, HRC, we saw much larger proportion of well-imputed exonic variants (gain 25.7% for MAC 10-30, 8.7% for 30-70 and 39.2% for 70-110.), showing the benefit of having both population-specific reference panel and the added extra information on exonic regions achieved by using the WES extension of the panel. Similar results for the non-coding variation are shown in **Supplementary Figure 1** for comparison.

In addition to the number of well-imputed variants, we calculated the false positive rate for each of the panels. This was done by focusing on the imputed genotypes of monomorphic sites that were seen in the original Human CoreExome genotype data. In total we identified 8,160 monomorphic sites in Finns in the chromosome 17. The monomorphic sites that were imputed as polymorphic (N = 898 with at least one heterozygous or minor allele homozygous genotype call with maximum posterior probability of > 0.9) were manually checked to be truly monomorphic using the genotyping array intensity data. We observed high false positive rates when using the global reference panels (4.0% for 1000G and 2.6% for HRC, **Figure 1C**) but not for the local reference panel. This result illustrates the difference between the Finnish genomes and more outbred populations with Finns having considerably less rare variants. When using global reference panels, many of these variants are falsely imputed to the target datasets.

The summary of the imputation of the exonic masked variation for the lowest allele count ranges (10-30, 30-70 and 70-110) can be seen in the **Figure 1D**). The number of masked variants present in the panel (panel sensitivity) was lowest in the 1000G panel, as was expected due to the low number of matching haplotypes, where 43% of the masked exonic variants were not present (presented in white in the **Figure 1D**). Highest sensitivity was seen for the HRC panel but when applying info cutoff of 0.7, the numbers of well-imputed variants were similar between HRC and local panel, but the accuracy of the imputation was best for the local reference panel, where we saw highest proportions of variants with concordance > 0.8 for all three MAC ranges. Similar results for masked variants including non-coding variants included are presented in the **Supplementary Figure 2**.

When combining two panels with partially overlapping variants, like low-coverage WGS and high-coverage WES, the panels need to be cross-imputed as has been suggested for example by the UK10K working group^6^. In IMPUTE2 this can be done simply by adding extra option to the command line. However, this option should be used with care. When using this method with a global panel and a population specific panel (e.g. with 1000G and low-pass WGS), the false positive rate of variants increased considerably (7.4%, **Figure 1C**). The false positive imputed genotypes also seemed to have high information scores with the combination panel; almost half of the false positive variants had info > 0.7. In case of a population specific panel and a global panel, the between panel imputation seems to introduce variation into the population specific panel, that does not really exist in that population, and as the number of “observed” alleles rises in the combined panel, the imputation method is more confident of the existence of that variation, therefore inflating the confidence as presented by the information scores. When applying info threshold of 0.9 instead of 0.7, we were able to lower the number of false positives (**Supplementary Table 1**) with the cost of filtering out also true positive variants, but were still left with 0.7% false positive rate for the combination panel of 1000G + low-pass WGS.

Based on these results, we conclude that the local reference panel outperforms the global imputation panels when imputing low frequency and rare variants, even when there is a considerable proportion of individuals from the same population in the global reference panel. Using a local panel not only provides largest number of successfully imputed variants, but also the lowest number of false positive rare and low frequency variants. In addition, it is clear that the existing large-scale WES data can be used to boost imputation quality of the exonic variation with very low allele frequencies. Combining the population-specific WGS panel with WES panel provides a powerful tool particularly for studies of the downstream health effects of low frequency and rare coding variants in large-scale association studies.

## Online Methods

### Study data set

Study data set comprises of 10,489 individuals from the National Finrisk study^12^. The individuals come from two separate genotyping projects. Individuals that were also in the reference data set were removed before the imputation. Sample-wise quality control included checks for heterozygosity (± 3sd from mean *F*), gender discrepancies and call-rate < 98%. Marker-wise QC included removal of indels, duplicates, markers with HWE *P*-value < 10^-6^ and markers with call rate < 95%. Genotyping algorithm used for both data sets was zCall^14^. The study data set was phased with ShapeIT2^15^ using default options and effective population size of 11,418.

### Finnish low-pass WGS reference data set

Sample quality control for the data was done at the Wellcome Trust Sanger Institute. SNPs with HWE *P*-value < 10^-5^ (99,191) were removed and the data was phased using ShapeIT2 using default options and effective population size of 11,418. There were 38,666 markers with discordant alleles compared to 1000 genomes September 2013 reference panel. Only polymorphic autosomal SNPs were included in the reference panel.

### Finnish WES reference data set

From the raw whole exome sequences variants were removed based on the following criteria: 1) Non-biallelic variants removed, 2) Genotypes with QC < 20 were set to missing, 3) SNPs with call rate < 95% were removed, and 4) Monomorphic markers were removed. In addition, samples that were also in the WGS panel (7) were excluded together with individuals whose genotyping rate was < 95% (43). After filtering steps, the reference panel contains 222,342 markers and 4,932 individuals. The data was phased using ShapeIT2 using default options and effective population size of 11,418.

### Imputation

The chromosome 17 of study data set was imputed using all reference panels described above after masking variants with MAF < 1% for studying imputation accuracy and number of variants that can be imputed with good quality. In the case of non HRC reference panels, IMPUTE^13^ version 2.3.1 was used with effective population size of 11,418 and Hidden Markov Model (HMM) states for imputation was set to the same value as the number of haplotypes in reference panels. HRC imputation was run with default settings using pre-phased study haplotypes.

### Concordance calculation

The variant concordance rate was calculated using imputed best-guess genotypes with posterior probability > 0.9 which is often the default in different software transforming probability based data into genotype calls. We used GTOOL (http://www.well.ox.ac.uk/~cfreeman/software/gwas/gtool.html) for the transformation. The concordance value for a variant for each of the panels was defined as the proportion of heterozygous and minor allele homozygous genotypes that matched to the original genotype data.

## Acknowledgements

This work was supported by the Academy of Finland (298149 to I.S.; 251217, 255847 and 285380 to S.R.; 251704 and 286500, 293404 to A.P.), the Academy of Finland Center of Excellence for Complex Disease Genetics of the Academy of Finland (grant nos. 213506 and 129680), the Wellcome Trust (WT089061, WT089062 and WT0980512), EU FP7 projects ENGAGE (201413 to A.P. & S.R.), BioSHaRE (261433 to A.P. and S.R.), the Finnish Foundation for Cardiovascular Research (V.S., A.P. and S.R.), the Sigrid Juselius Foundation (A.P. and S.R.), Biocentrum Helsinki (S.R.), the Nordic Information for Action eScience Center (NIASC)-a Nordic Center of Excellence financed by NordForsk (grant no. 62721 to A.P., & S.R.).

## References

1 McCarthy, S., et al. A reference panel of 64,976 haplotypes for genotype imputation. Nat. Genet. 48, 1279–1283 (2016)

2 Huang, L., et al. Genotype-imputation accuracy across worldwide human populations. Am. J. Hum. Genet. 84, 235–250 (2009)

3 1000 Genomes Project Consortium, et al. An integrated map of genetic variation from 1,092 human genomes. Nature 491, 56–65 (2012)

4 Gudbjartsson, D. F., et al. Large-scale whole-genome sequencing of the Icelandic population. Nat. Genet. 47, 435–444 (2015)

5 Genome of the Netherlands Consortium, Whole-genome sequence variation, population structure and demographic history of the Dutch population. Nat. Genet. 46, 818–825 (2014)

6 Huang, J., et al. Improved imputation of low-frequency and rare variants using the UK10K haplotype reference panel. Nat. Commun. 6, 8111 (2015)

7 Surakka, I., et al. Founder population-specific HapMap panel increases power in GWA studies through improved imputation accuracy and CNV tagging. Genome. Res. 20, 1344–1351 (2010)

8 Lek, M., et al. Analysis of protein-coding genetic variation in 60,706 humans. Nature 536, 285–291 (2016)

9 Jakkula, E., et al. The genome-wide patterns of variation expose significant substructure in a founder population. Am. J. Hum. Genet. 83, 787–794 (2008)

10 Lim, E. T., et al. Distribution and medical impact of loss-of-function variants in the Finnish founder population. PLoS Genet. 10, e1004494 (2014)

11 Mohlke, K. L., et al. Marker–marker linkage disequilibrium extends beyond 1 cM on chromosome 20 in Finns. Genome. Res. 11, 1221–1226 (2001)

12 Vartiainen, E., et al. Thirty-five-year trends in cardiovascular risk factors in Finland. Int. J. Epidemiol. 39, 504–518 (2010)

13 Howie, B. N., Donnelly, P. and Marchini, J. A flexible and accurate genotype imputation method for the next generation of genome-wide association studies. PLoS Genet. 5, e1000529 (2009)

14 Goldstein, J. I., et al. zCall: a rare variant caller for array-based genotyping: genetics and population analysis. Bioinformatics 28, 2543–2545 (2012)

15 Delaneau, O., Zagury, J. F. and Marchini, J. Improved whole chromosome phasing for disease and population genetic studies. Nat. Methods. 10, 5–6 (2013)

